# Endocytic Trafficking Promotes Vacuolar Enlargements for Fast Cell Expansion Rates in Plants

**DOI:** 10.1101/2021.11.29.470358

**Authors:** Kai Dünser, Maria Schöller, Christian Löfke, Nannan Xiao, Barbora Pařízková, Stanislav Melnik, Eva Stöger, Ondřej Novák, Jürgen Kleine-Vehn

## Abstract

The vacuole has a space-filling function, allowing a particularly rapid plant cell expansion with very little increase in cytosolic content (Löfke et al., 2015; Scheuring et al., 2016; Dünser et al., 2019). Despite its importance for cell size determination in plants, very little is known about the mechanisms that define vacuolar size. Here we show that the cellular and vacuolar size expansions are coordinated. By developing a pharmacological tool, we enabled the investigation of membrane delivery to the vacuole during cellular expansion. Counterintuitively, our data reveal that endocytic trafficking from the plasma membrane to the vacuole is enhanced in the course of rapid root cell expansion. While this “compromise” mechanism may theoretically at first decelerate cell surface enlargements, it fuels vacuolar expansion and, thereby, ensures the coordinated augmentation of vacuolar occupancy in dynamically expanding plant cells.

---

Animals and plants take many shapes and come in great diversity of sizes (Marshall et al., 2012). During cellular expansion surface area to intracellular volume ratio gets smaller and if cells grow beyond a critical limit, the surface of the plasma membrane may cease to accommodate cellular needs. Cells typically remain, hence, relatively small or eventually induce membrane furcation to enlarge the cellular surface. Compared to animals, plant cells can dramatically increase their size without the apparent need for surface enlargements. The vacuole is the biggest organelle in plants and its size correlates with and determines cell size in *Arabidopsis* (Owens and Poole, 1979; Berger et al., 1998; Löfke et al. 2015; Scheuring et al., 2016). Vacuolar size control is dynamic and vacuolar morphology is a reliable intracellular read-out for cellular expansion (Dünser et al., 2019). The size of the vacuole increases during cell expansion, occupying up to 90% of a cellular volume. Accordingly, the dynamic regulation of vacuolar size allows plant cells to expand with little increase of the cytosol (Dünser et al., 2019), maintaining a favorable cell surface to cytosol ratio during growth. Despite its anticipated importance for fast cellular expansion, very little is known about how cells scale and coordinate its vacuolar size during the highly dynamic process of cell enlargement (Dünser et al., 2015; Krüger and Schumacher, 2017). Actin- and myosin-dependent vacuolar constrictions define the vacuolar occupancy of the cell (Scheuring et al., 2016). On the other hand, vacuolar size could also relate to the control of vesicle trafficking, because the phytohormone auxin defines cell size in a soluble N-ethylmaleimide-sensitive-factor attachment receptor (SNARE)-dependent manner (Löfke et al., 2015). The vacuolar SNARE complex, consisting of R-SNARE VAMP711, Qa-SNARE SYP22, Qb-SNARE VTI11, as well as Qc-SNAREs SYP51 and SYP52, regulates hetero and homotypic membrane fusion at the tonoplast (Fujiwara et al., 2014). Auxin stabilizes vacuolar SNAREs in a posttranslational manner and *vti11* mutant vacuoles are resistant to the auxin effect on vacuoles, leading to auxin resistant root growth (Löfke et al., 2015).

Despite our molecular knowledge on vesicle trafficking towards the vacuole (Krüger and Schumacher, 2017), very little is known about the rate regulation of membrane delivery to the vacuole, its dynamics, and possible implication for cellular expansion. In this project we hypothesized that the interference with vacuolar SNAREs could aid us to visualize the rate and importance of membrane delivery to vacuole. Mutations in the vacuolar SNARE subunit *VTI11* leads to strongly affected, roundish, and partially fragmented vacuoles (Yano et al., 2003; Zheng et al., 2014; Löfke et al., 2015). Despite the strongly affected vacuolar morphology, the root length, as well as the size of fully elongated epidermal cells of untreated *vti11* mutants, were hardly distinguishable from wild type seedlings (Fig. 1 A - D). This finding seemingly questions the contribution of vacuolar SNARE-dependent membrane fusions at the tonoplast for cellular expansion. We, however, have proposed that the space-filling function (cellular occupancy) of the vacuole defines cell size determination (Dünser et al., 2019). Accordingly, we analyzed the cellular occupancy of the vacuole in the meristematic and elongation zone, using confocal z-stack imaging and 3-D renderings in epidermal atrichoblast cell files (Fig. 1 E). Although *vti11* mutant vacuoles are morphologically distinct from wild-type vacuoles (Yano et al., 2003; Zheng et al., 2014; Löfke et al., 2015), cellular occupancy of the vacuole in epidermal root cells was not distinguishable from wild-type (Fig. 1 F). This finding suggests that compensatory mechanisms, including the roundness of *vti11* mutant vacuoles, ensure the required cellular occupancy of the vacuole and, thereby, likely safeguards cell expansion.

**Figure 1.**
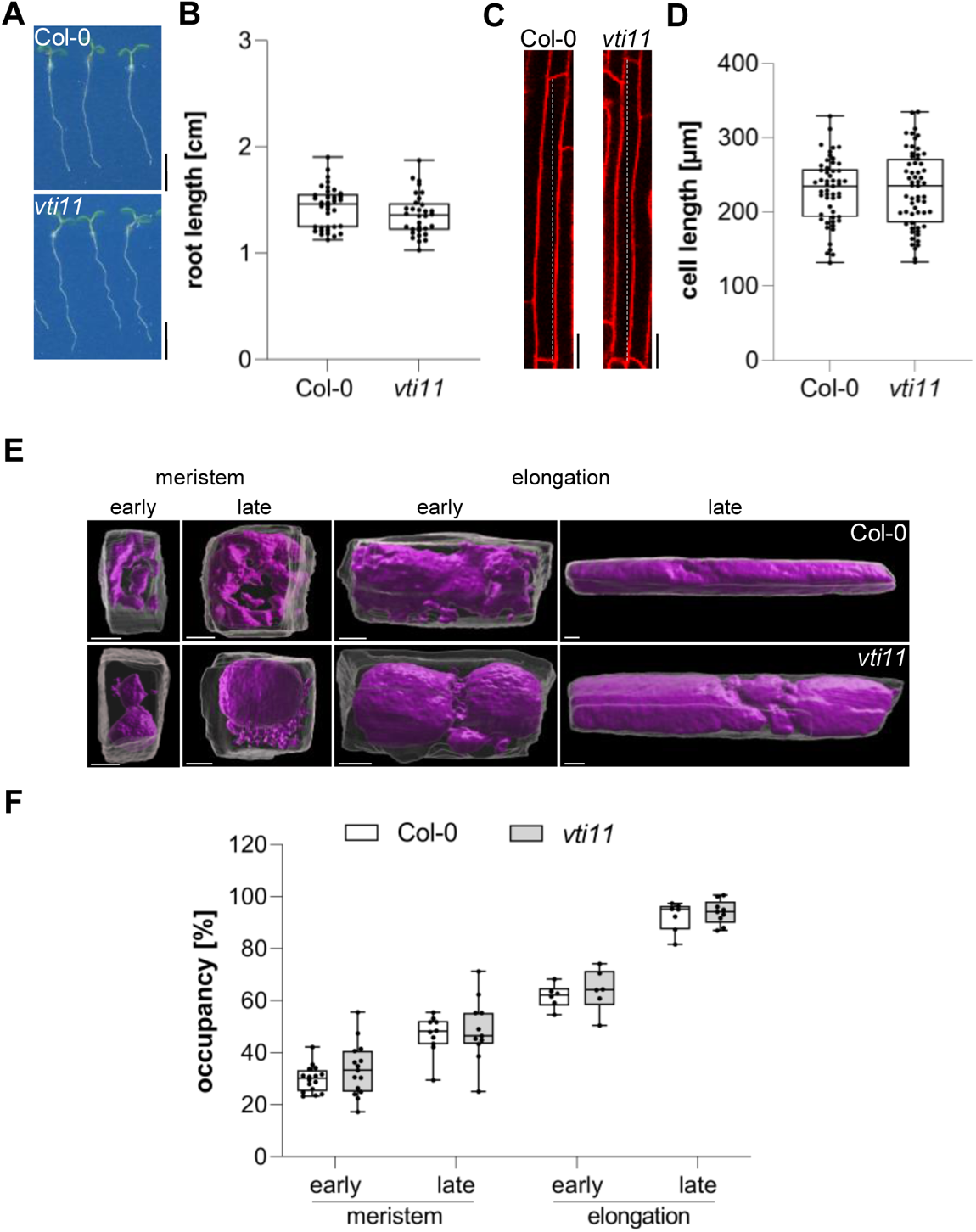
Vacuolar occupancy of the cell defines cell expansion. (A) Representative images (scale bars: 0.5 cm) and (B) quantification of the main root length of 6-day-old Col-0 (n = 36) and *vti11* seedlings (n = 32). Student’s t-test (ns). (C) Representative images (scale bars: 25 µm) of differentiated atrichoblast cells. Seedlings were stained with propidium iodide (PI) (red) for 30 min in liquid medium prior image acquisition. (D) Boxplots show cell length quantification of 6-day-old Col-0 (n = 53) and *vti11* (n = 59) seedlings. Student’s t-test (ns). (E) 3D reconstructions of PI-stained cell walls (grey) and BCECF-stained vacuoles (magenta) in the early and late meristem and in the early and late elongation zone. Scale bars: 5 µm. (F) Boxplots show vacuolar occupancy of cells in the defined zones (n = 7 – 16). Student’s t-test (ns). Boxplots: Box limits represent 25^th^ percentile and 75^th^ percentile; horizontal line represents median. Whiskers display min. to max. values. Data points are individual measured values. Representative experiments are shown.

*VTI11* and *VTI12* are able to substitute each other in their respective SNARE complexes with the *vti11 vti12* double mutants being lethal (Surpin et al., 2003). In order to overcome genetic redundancy as well as lethality in the crucial control of vesicle trafficking towards the vacuole, we conducted a small molecule screen with the aim to identify compounds impacting on SNARE-dependent delivery of vesicles to the vacuole. For our primary screen, we germinated seedlings constitutively expressing the vacuolar R-SNARE marker pUBQ10::YFP-VAMP711 in growth medium supplemented with bioactive chemicals from a library of 360 compounds, which likely affect cell expansion *in planta* (Drakakaki et al., 2011) (Fig. 2 A). We subsequently employed a fluorescence binocular to identify compounds that intensified the YFP-VAMP711 signal (Fig. 2 B). In a secondary, CLSM (confocal laser scanning microscopy)-based validation screen, we recorded vacuolar morphology changes in the samples treated with the compounds in comparison to solvent (control) treatments (Fig. 2 C, Suppl. Fig. 1 A). Thereby, we isolated 12 small molecules (Suppl. Fig. 1 A, B) that affected YFP-VAMP711 and vacuolar morphology (Suppl. Fig. 1 A), which we thus named <u>V</u>acuole <u>A</u>ffecting <u>C</u>ompounds (VACs) 1-12.

**Figure 2.**
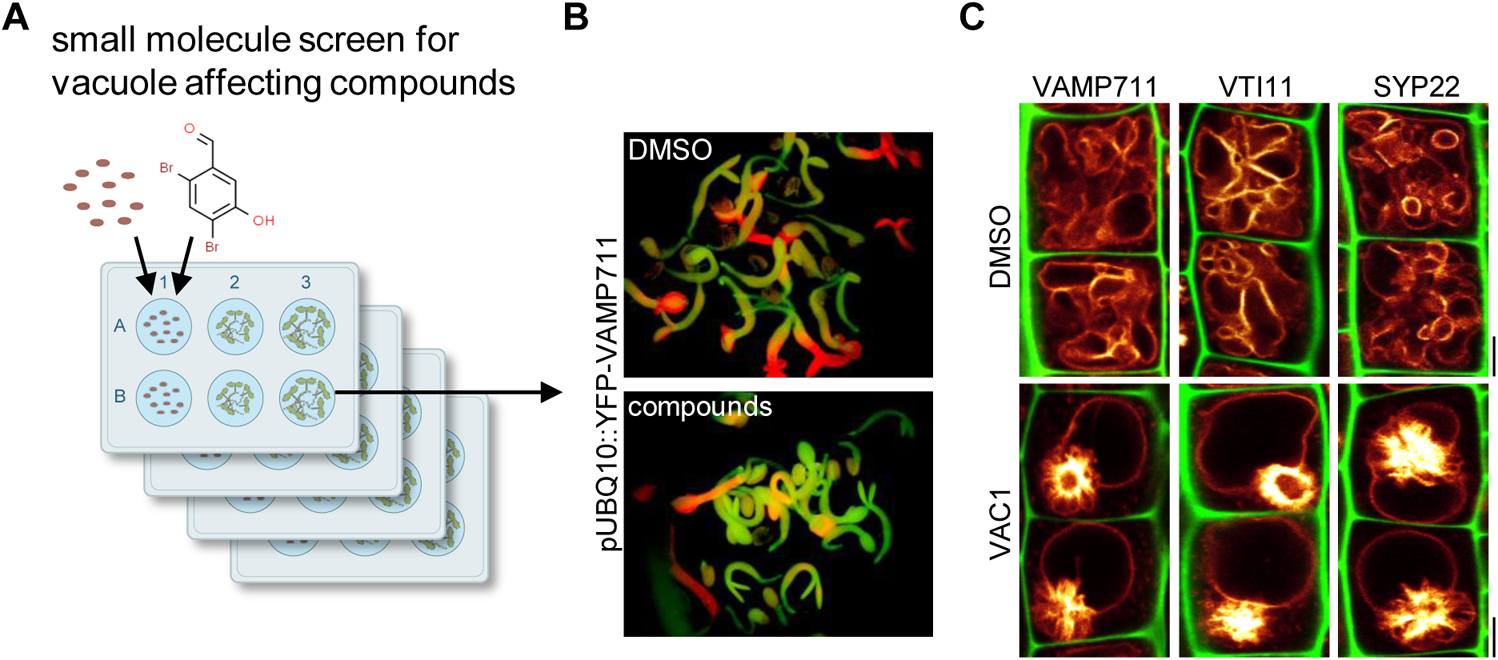
Bioactive small molecule screen identifies VAC1 as a vacuole affecting compound. (A) Schematic depiction of the small molecule screen workflow. pUBQ10::YFP-VAMP711 seeds were germinated in liquid medium containing solvent control DMSO or 360 small molecules from a library of bioactive compounds. (B) 4-day-old seedlings were then screened for intensified YFP-VAMP711 signal using a fluorescence binocular. (C) Representative images of the CLSM-based confirmation screen. Cell wall and vacuolar membrane in late meristematic atrichoblast cells were visualized with PI (green) and pUBQ10::YFP-VAMP711 (yellow), pSYP22::SYP22-GFP (yellow) or pUBQ10::pHGFP-VTI11 (yellow), respectively. Scale bars: 5 µm.

Especially VAC1 (N-[(2,4-Dibromo-5-hydroxyphenyl)methylideneamino]-2-phenylacetamide) had a striking effect on vacuolar morphology (Suppl. Fig. 1 A) and induced ectopic accumulation of vacuolar SNARE marker YFP-VAMP711 adjacent to the main vacuole (Fig. 2 C). Due to the possibly hydrolysable acylhydrazone bond in VAC1, we wondered whether VAC1 could be potentially metabolised in planta into some of its sub-structures or other metabolic products. VAC1 can be synthesized from (and potentially decompose back into) 2,4-dibromo-5-hydroxybenzaldehyde and phenylacetic acid hydrazide, but the application of these substances (here referred to as precursors) did not cause any alterations to vacuolar morphology (Suppl. Fig. 2 A). We also analyzed the potential VAC1 conversion to phenylacetic acid (PAA), because it is a naturally occurring compound with auxin activity (Abe et al., 1974). The ultra-high performance liquid chromatography–selected ion recording–mass spectrometry (UHPLC–SIR–MS) analysis detected VAC1, but not PAA in VAC1-treated seedlings (Suppl. Fig. 2 B-D). We conclude that VAC1 is not detectably converted to PAA under our conditions, suggesting an alternative mode of action. On the other hand, in VAC1-treated samples the UHPLC-UV-MS analysis revealed three additional peaks for potential VAC1 metabolites (M1-M3) with a retention time of 4.37 min, 5.26 min and 5.39 min for M1, M2 and M3, respectively (Suppl. Fig 2 E-F). High-resolution mass spectrometry (HRMS) was then employed to identify these metabolites based on exact mass determination and characteristic distribution of bromine isotopes (Suppl. Fig. 2 G). Metabolite M1 was predicted to be the hydroxylated form (VAC-OH, Rt 4.37 min, [M-H^−^]: 424.9129 Da, calculated elemental composition C_15_H_11_Br_2_N_2_O_3_) and the other two derivatives relate to glycosylated forms. The detected compound M2 likely represented VAC1-3-O-glucoside (VAC1-Glc, Rt 5.26 min, [M-H^−^]: 570.9716 Da, C_21_H_22_Br_2_N_2_O_7_) and M3 was identified as VAC1-3-O-(6-O-acetylglucoside) (VAC1-AcGlc, Rt 5.39 min, C_23_H_24_Br_2_N_2_O_8_, [M-H^−^]: 612.9805 Da) (Suppl. Fig. 2 G). Using UHPLC–UV (λ_max_ = 291 nm for all compounds), we then revealed VAC1-AcGlc (56.3 ± 0.8 %) as the most abundant metabolite, followed by VAC1-Glc (31.3 ± 0.4 %) and VAC1 (12.4 ± 0.6 %), while the UV signal of hydroxylated VAC1-OH (< 0.1 %) was close to the detection limit of the photodiode array detector used (Suppl. Fig. 2 H). This set of data shows that a substantial amount of VAC1 undergoes glycosylation *in planta*, which could affect its activity in long-term experiments.

Next, we used structurally related derivatives of VAC1, termed VAC1-analogue (VAC1A), to further address whether the entity of VAC1 is required for its effects on vacuolar morphology (Suppl. Fig. 3 A). At the concentration and treatment time used for the subcellular effect of VAC1 (10 µM for 2.5 h), solely VAC1A4, which displays an additional hydroxy group, induced severe vacuolar morphology defects (Suppl. Fig. 3 B). The cellular effects of VAC1A4 were, however, visually distinct from VAC1. While short term (2.5 h) VAC1A4 treated cells appeared viable (based on the intact extracellular propidium iodide staining) (Suppl. Fig. 3 B), the main root growth did not resume after wash-out (Suppl. Fig. 3 C), indicating some degree of toxicity. On the other hand, interference with either the phenyl-group or the functional groups of the 2,4-dibromo-5-hydroxybenzaldehyde reduced the activity of VAC1 (Suppl. Fig. 3 B). At higher concentrations (25 µM and 50 µM), VAC1A2 and VAC1A5 also impacted to some degree on vacuolar morphology, while VAC1A1 and VAC1A3 appeared largely inactive in regards to vacuolar morphology (Suppl. Fig. 3 B). These results imply that the entity of VAC1 (phenyl-as well as the benzaldehyde rings) contributes to its effect on vacuolar morphology.

**Figure 3.**
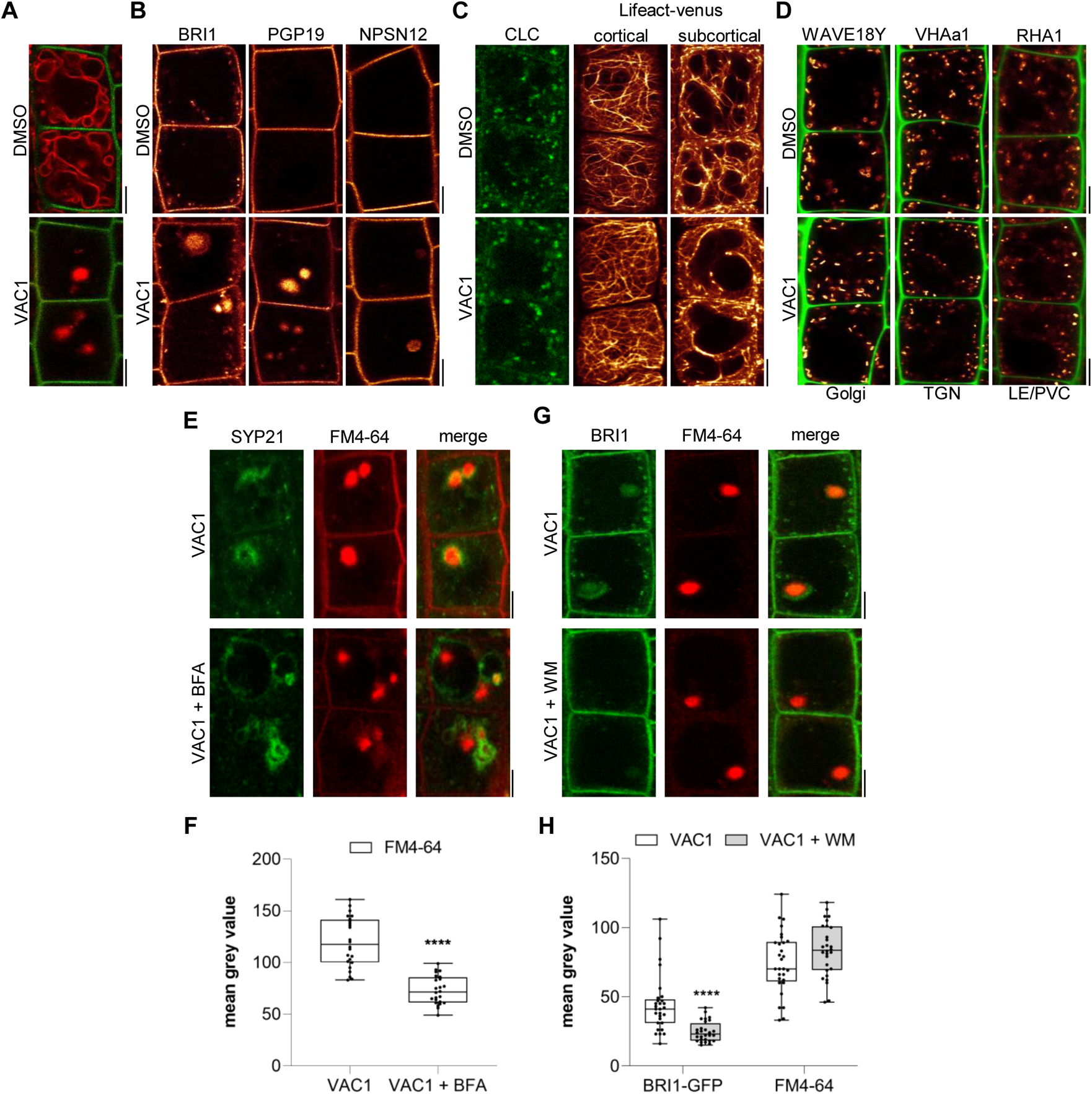
VAC1 interferes with vesicle trafficking to the vacuole. (A) Tonoplast and plasma membrane of late meristematic atrichoblast cells was visualized with FM4-64 (red) and pUBQ10::NPSN12-YFP (green), respectively. 6-day-old seedlings were pre-treated with DMSO or 10 µM VAC1 for 30 min., then pulse-stained with 4 µM FM4-64 for 5 min and subsequently transferred to liquid medium containing DMSO or 10 µM VAC1 for 3 h. Scale bars: 5 µm. (B) Representative images of 6-day-old seedlings of plasma membrane marker lines pBRI1::BRI1-GFP (yellow), pPGP19::PGP-19-GFP (yellow) and pUBQ10::NPSN12-YFP (yellow). Seedlings were treated with DMSO or 10 µM VAC1 for 2.5 h in liquid medium. Scale bars: 5 µm. (C) Representative images of 6-day-old seedlings of CLC-GFP (green) and Lifeact-venus (yellow) marker lines. Seedlings were treated with DMSO or 10 µM VAC1 for 2.5 h in liquid medium. Scale bars: 5 µm. (D) Representative images of 6-day-old WAVE18Y (pUBQ10::GOT1-YFP) (yellow), pVHAa1::VHAa1-GFP (yellow) and WAVE7Y (pUBQ10::RABF2a-YFP) (yellow) treated with DMSO or 10 µM VAC1 for 2.5 h in liquid medium. Cell walls were counter-stained with PI. Scale bars: 5 µm. (E) 6-day-old p35S::SYP21-GFP (green) seedlings were pulse stained with 4 µM FM4-64 (red) for 5 min and subsequently pre-treated with 25 µM BFA for 30 min before adding 10 µM VAC1 or solvent control for another 60 min. The treatments were conducted in liquid medium. Scale bars: 5 µm. (F) Boxplots show mean grey values of FM4-64 fluorescence signal in VAC1 bodies in VAC1 (n = 24) or VAC1 + BFA (n = 24) treatments. Student’s t-test (****P < 0.0001). (G) 6-day-old pBRI1::BRI1-GFP (green) seedlings were pulse stained with 4 µM FM4-64 (red) for 5 min and subsequently pre-treated with 33 µM WM for 30 min before adding 10 µM VAC1 or solvent control for another 60 min The treatments were conducted in liquid medium. Scale bars: 5 µm. (H) Boxplots show mean grey values of FM4-64 or BRI1-GFP fluorescence signal in VAC1 bodies in VAC1 (n = 29 for FM4-64, n = 29 for BRI1-GFP) or VAC1 + WM (n = 28 for FM4-64, n = 28 for BRI1-GFP) treatments. Student’s t-test (****P < 0.0001). Boxplots: Box limits represent 25^th^ percentile and 75^th^ percentile; horizontal line represents median. Whiskers display min. to max. values. Data points are individual measured values. Representative experiments are shown.

Next, we addressed the subcellular effects of VAC1 on the endomembrane system. When compared to YFP-VAMP711, we observed similar VAC1 effects on vacuolar morphology and protein mislocalization in *pUBQ10::pHGFP-VTI11*- and *pSYP22::SYP22-GFP*-expressing seedlings (Fig. 2 C). This finding suggests that VAC1 generally interferes with the localization of the SNARE complex. To further address whether this VAC1 effect disrupts vesicle trafficking to the vacuole, we next employed the styryl dye FM4-64 (Scheuring et al., 2015). The endocytic tracer FM4-64 initially labels the plasma membrane and is rapidly internalized following the endocytic pathway, eventually staining the vacuolar membrane (Scheuring et al., 2015). Tonoplast labelling with FM4-64 was observed after 3 hours following the pulse staining in mock-treated seedlings (Fig. 3 A). Strikingly, staining of the vacuolar membrane was absent in VAC1 treated samples and instead FM4-64 accumulated in VAC1 bodies (Fig. 3 A). This piece of data demonstrates that VAC1 treatments prevent the fusion of FM4-64 positive endocytic vesicles to the central vacuole. Next, we inspected plasma membrane resident proteins BRI1-GFP, PGP19-GFP and NPSN12-GFP, expecting a fraction of these proteins transiting to the vacuole for lytic degradation (Kleine-Vehn et al., 2008). When compared to the ectopic FM4-64 accumulations, the examined lines showed comparably weak, but clearly detectable VAC1-induced aggregation of the fluorescently tagged proteins, which were absent in the solvent control treatments (Fig. 3 B). This finding corroborates our assumption that VAC1 interferes with vesicle trafficking to the vacuole. Moreover, we conclude that VAC1 does not disrupt endocytic trafficking. In agreement, VAC1 did not visibly affect clathrin light chain marker CLC-GFP and F-actin marker Lifeact-venus (Fig. 3 C), suggesting that basic cellular functions, such as endocytosis and the actin cytoskeleton likely remain unaffected by VAC1 treatments. In agreement, also early and late endosome marker lines, such as GOT1-YFP, VHA-a1-GFP and RABF2A-YFP, were not visibly affected upon VAC1 administration (Fig. 3 D). This set of data suggests that early vesicle sorting towards the vacuole is likely not disturbed.

Next, we used Brefeldin A (BFA), which is a specific inhibitor of ARF-GEFs (guanine-nucleotide exchange factors for ADP-ribosylation factor GTPases), to block endocytic trafficking at the level of the trans-Golgi network (TGN). FM4-64 accumulation into VAC1 bodies (labelled by SYP21-GFP) was significantly reduced in BFA co-treatments (Fig. 3 E, F), suggesting that VAC1 affects vesicle trafficking downstream of the TGN.

The TGN functions as a sorting hub, selectively allowing membranes to transit via the multivesicular bodies (MVBs) or MVB independent trafficking routes to the vacuole. We, subsequently, used the PI3- and PI4-kinase inhibitor wortmannin (WM) to pharmacologically interfere with the function of the MVBs (also called pre-vacuolar compartments (PVCs) in plants) (Kleine-Vehn et al., 2008) (Fig. 3 G, H). WM application did not affect the FM4-64 signal intensities in VAC1 bodies, which implies MVB/PVC-independent membrane trafficking towards the VAC1 bodies (Fig. 3 G, H). In contrast, signal intensities of BRI1-GFP in the VAC1 bodies were significantly reduced in WM and VAC1 co-treatments when compared to VAC1 treatments alone (Fig. 3 G, H). These results demonstrate that VAC1 interferes with processes downstream of MVB/PVC-dependent, but also MVB/PVC-independent, vesicle trafficking routes to the vacuole.

To further decipher the VAC1 effect, we next addressed its dependency on the CLASS C CORE VACUOLE/ENDOSOME TETHERING (CORVET) and HOMOTYPIC FUSION AND PROTEIN SORTING (HOPS) machineries (Takemoto et al., 2018), because these membrane tethering complexes supply the vacuolar SNAREs with heterotypic (endosomal derived) and homotypic (vacuole derived) membranes, respectively. The dexamethasone (DEX)-inducible amiRNA lines that target either the CORVET-specific subunit VPS3 or the HOPS-specific subunit VPS39 enable the selective repression of the CORVET and HOPS complex, respectively (Takemoto et al., 2018). The transcriptional repression of CORVET or HOPS inhibits root growth (Fig. 4 A, B and Suppl. Fig. 4 A, B). In agreement, seedlings germinated on VAC1 containing media also displayed reduced main root growth (Fig 4 A, B and Fig. 4 F, G), suggesting that vesicle trafficking to the vacuole contributes to root growth. Homotypic membrane fusion unlikely contributes to the VAC1 effect, because VPS39 depleted roots remained fully sensitive to VAC1 (Suppl. Fig. 4 A-C). In contrast, the repression of CORVET function induced a partially resistant root growth to VAC1 (Fig. 4 A-C), suggesting that defects in heterotypic vesicle trafficking towards the vacuole contribute to the VAC1-dependent interference with root growth. To visualize vacuolar morphology and subcellular VAC1 effect after CORVET repression, we used the YFP-VAMP711 marker crossed to the dexamethasone (DEX)-inducible VPS3 amiRNA line (Takemoto et al., 2018). The conditional knockdown of VPS3 had no visible effect on the distribution of *pUBQ10::YFP-VAMP711* and the VPS3 deprived cells remained sensitive to VAC1-induced aberrations in VAMP711 localization (Fig. 4 D). This set of data suggests that CORVET-dependent vesicle tethering functions upstream of VAC1-sensitive processes.

**Figure 4.**
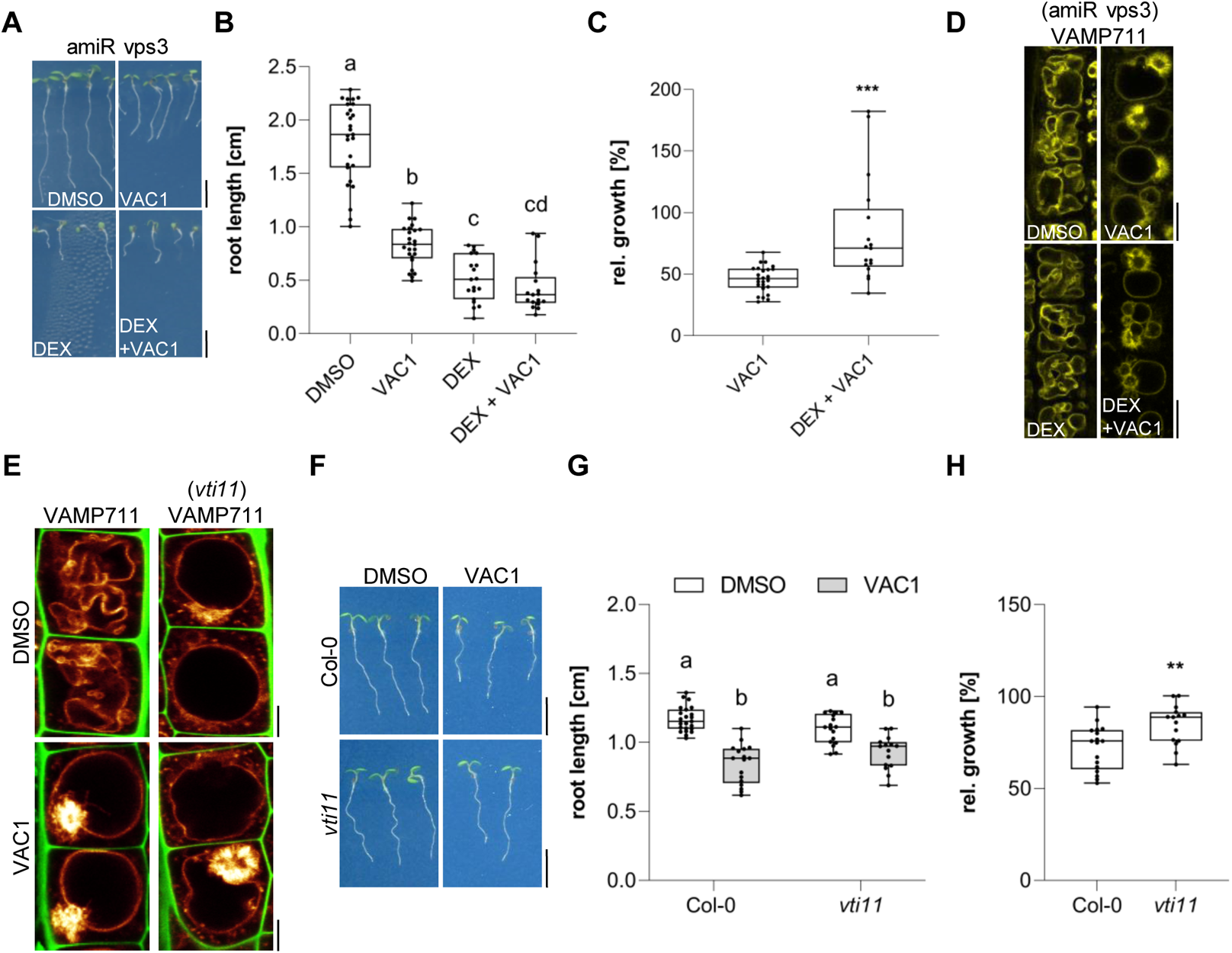
VAC1 specifically interferes with vacuolar SNARE-dependent vesicle fusion to the tonoplast. (A) Representative images (scale bars: 0.5 cm) and (B) quantification of main root length of 7-day-old amiR Vps3 seedlings germinated on solvent control medium (DMSO, n =27), 10 µM VAC1 (n = 25), 30 µM Dexamethasone (DEX, n = 19) or 10 µM VAC1 and 30 µM DEX (n =17). One-way ANOVA with Tukeýs multiple comparisons test (b: P < 0.0001, c: P < 0.0001, d to a and b: P < 0.0001). (C) Boxplots show relative growth of VAC1 samples (compared to DMSO) and DEX + VAC1 samples (compared to DEX). Student’s t-test (***P = 0.0002). (D) Representative images of late meristematic atrichoblast cells in *amiR Vps3 x pUBQ10::YFP-VAMP711* seedlings. Seedlings were grown for 4 days, then transferred to plates containing solvent control (DMSO) or 30 µM DEX for another 3 days und subsequently treated for 2.5 h with DMSO, 30 µM DEX, 10 µM VAC1 or VAC1 + DEX, respectively, in liquid medium. Scale bars: 5 µm. (E) Representative images of late meristematic atrichoblast cells of pUBQ10::YFP-VAMP711 in wild-type and *vti11* background. PI (green) and *pUBQ10::YFP-VAMP711* (yellow) depict cell wall and vacuolar membrane, respectively. 6-day-old seedlings were treated with solvent control (DMSO) or 10 µM VAC1 for 2.5 h in liquid medium. Scale bars: 5 µm. (F) Representative images (scale bars: 0.5 cm) and (G) boxplots showing main root length of 6-day-old Col-0 and *vti11* seedlings grown on solvent control (DMSO) (n = 23 for Col-0, n = 17 for *vti11*) or 20 µM VAC1 (n = 17 for Col-0, n = 15 for *vti11*) plates. 2way ANOVA indicated interaction (P = 0.0053). Statistical significance was determined by one-way ANOVA with Tukeýs multiple comparisons test (b: P ≤ 0.0006). (H) Boxplots show relative growth of Col-0 and *vti11* on 20 µM VAC1 (compared to DMSO). Student’s t-test (**P = 0.0051). Boxplots: Box limits represent 25^th^ percentile and 75^th^ percentile; horizontal line represents median. Whiskers display min. to max. values. Data points are individual measured values. Representative experiments are shown.

We, accordingly, propose that VAC1 primarily interferes with vesicle fusion at the tonoplast downstream of heterotypic vesicle tethering complex. In agreement, VAC1 treated wild type cells remarkably resembled the vacuolar morphology of untreated *vti11* mutants (Fig. 4 E) and, hence, we subsequently addressed the contribution of vacuolar SNARE *VTI11* to the VAC1 effect. We concluded that VAC1 attenuates root growth rates in a VTI11-dependent manner, because when compared to wild type seedlings the *vti11* mutant roots were resistant to VAC1 (Fig. 4 F-H). On the other hand, VAC1 application still induced alterations in VAMP711 localization in the *vti11* mutant background (Fig. 4 E). We accordingly conclude that VAC1 interferes with SNARE-dependent membrane delivery to the vacuole, but that VTI11 is not the sole or main target of VAC1, assuming other molecular targets and/or functional redundancy at the level of vacuolar SNAREs.

Considering that VAC1 interferes with vacuolar SNARE-dependent vesicle trafficking to the vacuole, this pharmacological tool hence allowed us to address the importance of vesicle trafficking towards the vacuole in elongating cells. We subsequently used VAC1 to visualize the rate of endocytic trafficking to the vacuole in early and late meristematic as well as early and late elongation zones. We used FM4-64 for pulse labelling of the plasma membranes (Suppl. Fig. 5 A and B) and observed that elongating cells displayed higher FM4-64 uptake into VAC1 bodies when compared to meristematic cells (Fig. 5 A and B). To address if this finding implies higher degree of endocytic membrane delivery into VAC1 bodies during cellular elongation, we similarly used BFA to block endocytic membranes already at the level of the TGN (Kleine-Vehn et al., 2008). In line with the results from VAC1 treatments, BFA bodies also showed greater signal intensities in cells of the elongation zone when compared to cells of the meristem (Fig. 5 C, D). In addition, we performed whole-cell 3-D reconstructions of early and late meristematic cells. While displaying a doubling of cell surface area, the volume of endocytic FM4-64 accumulation in BFA bodies quadrupled in late meristematic cells compared to early meristematic cells (Fig. 5 E, F). We, hence, propose that enhanced endocytic trafficking correlates with elongation.

**Figure 5.**
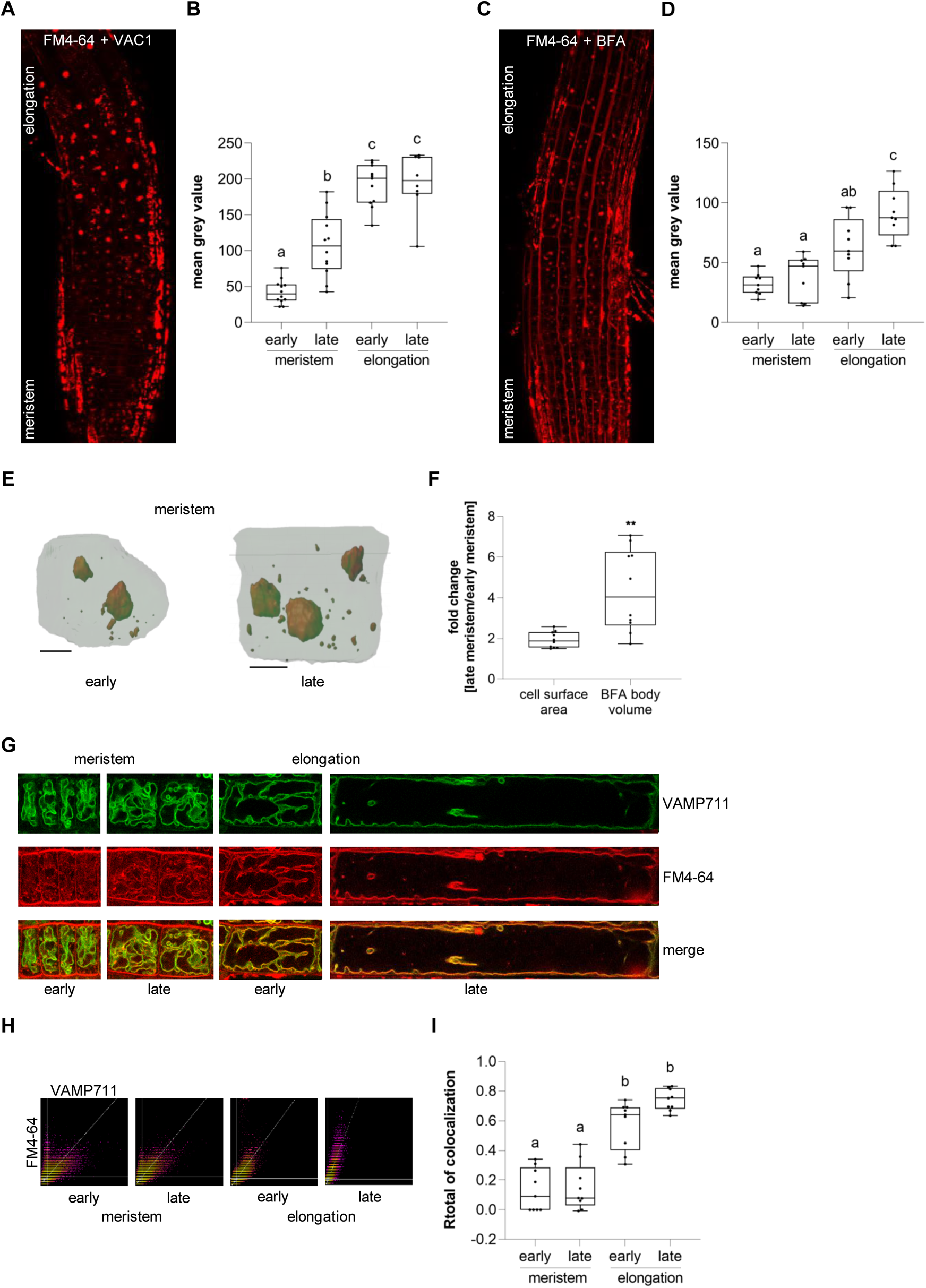
Endocytic trafficking is enhanced at the onset of cellular elongation. (A) Representative maximum z-projection (scale bar: 50 µm) of 6-day-old Col-0 roots. Seedlings were pre-treated with 10 µM VAC1 for 30 min before staining in 4 µM FM4-64 (red) for 30 min Seedlings were then de-stained for 1.5 h and subsequently treated with 10 µM VAC1 for 3 h before image acquisition. The treatments and the staining were conducted in liquid medium. (B) Boxplots showing mean grey values of FM4-64 fluorescence signal in VAC1-bodies in early (n = 12) and late (n = 12) meristematic cells as well as in the early (n = 11) and late (n = 8) elongation zone. One-way ANOVA with Tukeýs multiple comparisons test (b: P = 0.0002, c: P < 0.0001). (C) Representative maximum z-projection (scale bar. 50 µm) of 6-day-old Col-0 roots. Seedlings were pulse-stained in 4 µM FM4-64 (red) for 5 min and subsequently transferred to 25 µM BFA for 30 min prior to image acquisition. Staining and treatment was done in liquid medium. (D) Boxplots showing mean grey values of FM4-64 fluorescence signal in BFA-bodies in early (n = 9) and late (n = 9) meristematic cells as well as in the early (n = 9) and late (n = 9) elongation zone. One-way ANOVA with Tukeýs multiple comparisons test (b to a (early meristem): P = 0.0123, c to a: P < 0.0001, c to b: P = 0.0178). (E) 3D reconstructions of plasma membrane (pUBQ10::NPSN12-YFP, green) and BFA bodies (red) in early (left) and late (right) meristematic cells. 6-day-old pUBQ10::NPSN12-YFP seedlings were pulse-stained with 4 µM FM4-64 for 5 min and subsequently transferred to 50 µM BFA for 2 h prior image acquisition. Staining and treatment was done in liquid medium. Scale bars: 5 µm. (F) Boxplots depict fold change of cell surface area (n = 10) and BFA body volume (n =10) in late meristematic cells when compared to early meristematic cells. (G) Representative images of early and late meristematic cells and cells of the early and late elongation zone. 6-day-old pUBQ10::YFP-VAMP711 seedlings were pulse-stained with 4 µM FM4-64 for 5 min and subsequently de-stained in liquid medium for 3 h before image acquisition. Overlay of YFP-VAMP711 (green, upper panel) and FM4-64 (red, middle panel) fluorescence signals is shown in the lower panel. Scale bar: 10 µm. Note: Images are assembled on white background. (H) Representative scatter plots depict co-localization of YFP-VAMP711 and FM4-64 in cells of the early and late meristem and in the early and late elongation zone. (I) Boxplots show Rtotal of co-localization between YFP-VAMP711 and FM4-64 in early (n = 9) and late (n = 9) meristematic cells and in the early (n = 9) and late (n = 9) elongation zone. Boxplots: Box limits represent 25^th^ percentile and 75^th^ percentile; horizontal line represents median. Whiskers display min. to max. values. Data points are individual measured values. Representative experiments are shown.

Next, we tested whether endocytic trafficking remains enhanced during cellular elongation when we do not apply vesicle trafficking inhibitors. To this end, we quantified the co-localization of FM4-64 with the tonoplast marker YFP-VAMP711 in meristematic and elongating cells (Fig. 5 G). Three hours after the pulse staining of FM4-64 early meristematic cells displayed more plasma membrane labeling when compared to late meristematic and elongating cells (Fig. 5 G). Moreover, the co-localization between FM4-64 and YFP-VAMP711 at the vacuolar membrane was significantly lower in the meristematic zone when compared to the elongation zone (Fig. 5 H, I). Taken together, these results indicate that endocytic membrane sorting towards the vacuole undergoes significant rate changes during cellular elongation.

Based on our data, it is conceivable that endocytic trafficking may impact cell expansion via its contribution to vacuole size. To approach this, we initially quantified cell and vacuole surface areas of (early and late) meristematic and (early and late) elongating cells using 3D reconstructions. We found that the area increase of both cell and vacuole surface is remarkably similar during the course of cell expansion (Fig. 6 A (left panel), B). This result indicates that intensified membrane delivery is not only required at the plasma membrane, but similarly occurs at the tonoplast during cellular growth. Accordingly, not only actin- and myosin-dependent vacuolar unfolding (Scheuring et al., 2016), but also membrane delivery contributes to vacuolar size control. Our data, moreover, propose that cell and vacuolar enlargements are coordinated during rapid cellular expansion. To address this coordinative mechanism, we applied VAC1 and employed whole-cell 3D reconstructions (Fig. 6 A (right panel)). Within only 2.5 hours, VAC1 induced an already detectable imbalance of cell and tonoplast surface size (Fig. 6 C).

**Figure 6.**
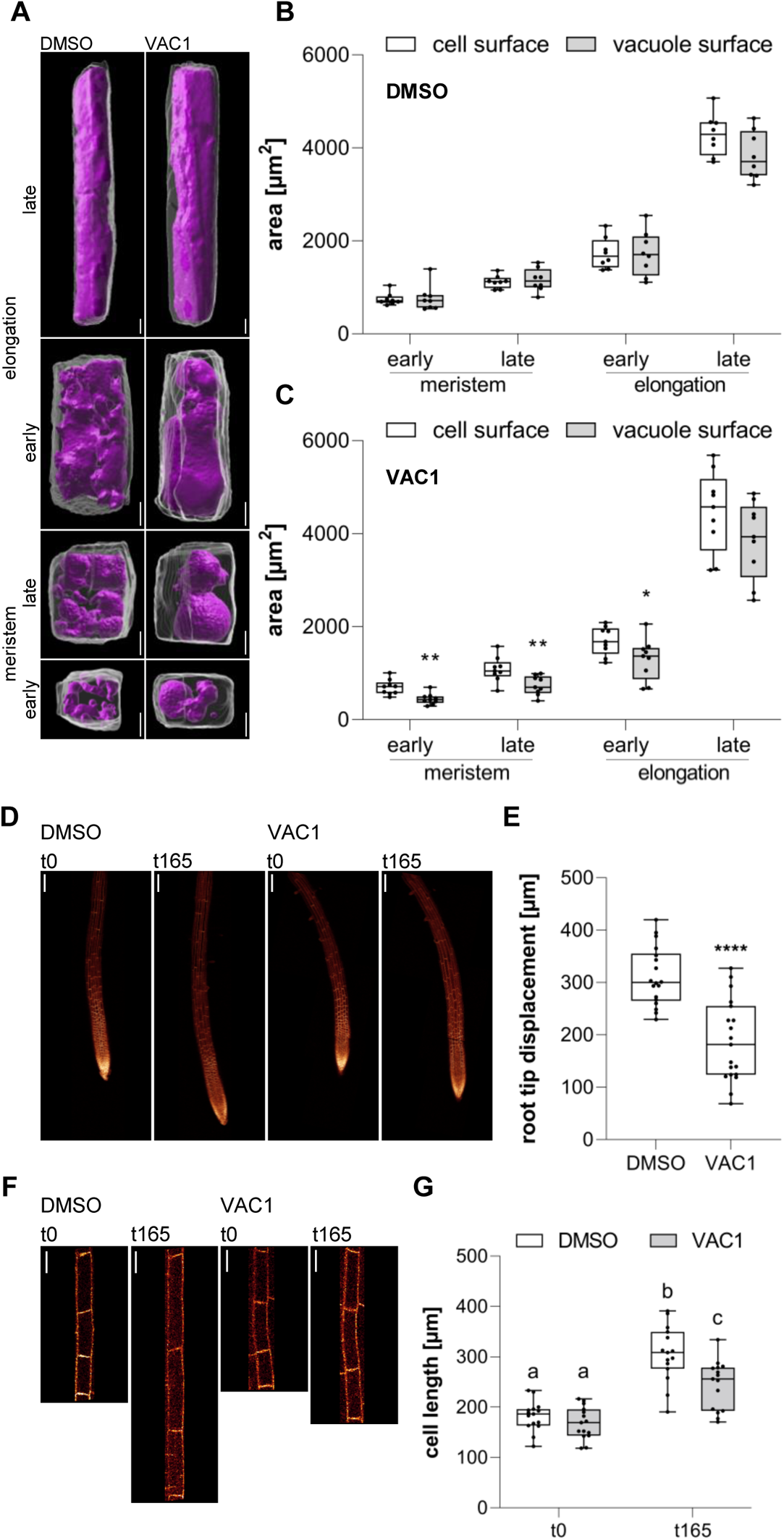
Coordinated surface increase at PM and tonoplast during cell elongation. (A) 3D reconstructions of propidium iodide (PI)-stained cell walls (grey) and BCECF-stained vacuoles (magenta) in the early and late meristem and in the early and late elongation zone. Scale bars: 5 µm. (B) and (C) Boxplots show cell and vacuole surface areas in the defined zones and indicated treatments. 6-day-old Col-0 seedlings were treated with solvent control (DMSO, n = 8) or 10 µM VAC1 (n = 9) for 2.5 h in liquid medium. Student’s t-test (early meristem: **P = 0.0018, late meristem: **P = 0.0073, early elongation: *P = 0.0423). (D) Representative images (scale bars: 100 µm) of maximum z-projections of 6-day-old pUBQ10::NPSN12-YFP (yellow) seedlings. Seedlings were mounted on agar blocks containing solvent control (DMSO) or 20 µM VAC1 in chamber slides. t0 represents the first and t165 the last image acquisition time point, given in minutes after mounting. Note: Images are assembled on black background. (E) Boxplots show root tip displacement of DMSO (n = 18) and VAC1 (n = 19) treated samples at the end of the 165 min time frame. Student’s t-test (****P < 0.0001). (F) Representative images (scale bars: 25 µm) of maximum z-projections of 6-day-old pUBQ10::NPSN12-YFP (yellow) seedlings. Seedlings were mounted on agar blocks containing solvent control (DMSO) or 20 µM VAC1 in chamber slides. t0 represents the first and t165 the last image acquisition time point, given in minutes after mounting. Note: Images are assembled on black background. (G) Boxplots show cell lengths of DMSO (n = 15) and VAC1 (n = 15) treated samples at the beginning and the end of the 165 min time frame. One-way ANOVA with Tukeýs multiple comparisons test (b to a: P < 0.0001, b to c: P = 0.0013, c to a: P ≤ 0.0012).

In summary, our set of data demonstrates (I) enhanced endocytic trafficking to the vacuole during cellular elongation and (II) that VAC1-reliant interference with SNARE-dependent vesicle fusion at the tonoplast impairs the coordination of plasma membrane and tonoplast surface area increase during cell enlargements.

Our data propose that the dynamic regulation of endocytic trafficking is contributing to the vacuole size increase, which could have an impact on rapid cell expansion rates. To address this, we used VAC1 to temporally block the vesicle fusion to the tonoplast during cell expansion. Accordingly, we mounted Col-0 seedlings in VAC1 or solvent (control) containing slide chambers and immediately inspected root growth under these conditions using confocal microscopy. VAC1-induced without a further delay a slower root growth (tip displacement) (Fig. 6 D, E), which correlated with reduced cellular expansion rates (Fig. 6 F, G) when compared to the solvent control. Accordingly, we conclude that vesicle trafficking to the vacuole is required for fast cellular expansion rates.

Here, we isolated and characterized VAC1, which interferes with the final steps of membrane delivery to the tonoplast. The molecular target of VAC1 is unknown, but the structure-activity assessment suggests that the entirety of the compound is contributing to its activity, possibly insulating a complex mode of VAC1 action. Our data suggest that VAC1 disrupts heterotypic vesicle delivery to the tonoplast. It is tempting to speculate that VAC1 interferes with possibly several subunits of the vacuolar SNARE complex or yet to be identified accessory proteins. Accordingly, in-depth characterization of the mode of VAC1 action could reveal intermolecular mechanisms of vesicle fusion at the tonoplast.

We used VAC1 to visualize the rate of vesicle trafficking towards the vacuole and thereby revealed intense reprogramming of endocytic membrane trafficking to the vacuole. Mechanistically, we show that endocytic trafficking is accelerated at the onset of elongation, suggesting that plasma membrane derived vesicles at least in part fuel the size increase of the vacuole. The development of this pharmacological tool allowed us to reveal the importance of vacuolar surface increase for cell expansion and overall root growth. The enhanced internalization of plasma membranes during cell expansion is somewhat counterintuitive, because it slows down the theoretically possible cell surface increase. On the other hand, this compromise model of vesicle trafficking based enlargements of plant cells allows the steady increase in vacuolar occupancy of the cell, which enables growth with little de novo synthesis of cytosolic components. This coordinated membrane flow mechanism appears to synchronize cell and vacuolar surface increase, which is likely the reason why plant cells excel in the speed of cellular expansion. It remains to be seen how plant cells molecularly reprogram and monitor the rate of endocytic trafficking for ensuring fast cellular elongation.

## Material and methods

### Plant material and growth conditions

Experiments were carried out in *A. thaliana* (Col-0 ecotype). The following plant lines were described previously: *vti11* (zizag) (Yano et al., 2003), *pUBQ10::VAMP711-YFP* (WAVE9Y) (Geldner et al, 2009), *pUBQ10::pHGFP-VTI11* (Takemoto et al., 2018), *pSYP22::SYP22-GFP* in *syp22* background (Uemura et al., 2010), *pBRI1::BRI1-GFP* (Kleine-Vehn et al., 2008), *pPGP19::PGP19-GFP* (Dhonukshe et al., 2008), *pUBQ10::NPSN12-YFP* (WAVE131Y) (Geldner et al., 2009), *CLC-GFP* (Ito et al., 2012), *p35S::Lifeact-venus* (Era et al., 2009), *pUBQ10::GOT1-YFP* (WAVE18Y) (Geldner et al., 2009), *pVHAa1::VHAa1-GFP* (Dettmer et al., 2006), *pUBQ10::RABF2a-YFP* (WAVE7Y) (Geldner et al., 2009), *p35S::SYP21-GFP* (Robert et al., 2008), *amiR vps3* (Takemoto et al., 2018), *amiR vps39* (Takemoto et al., 2018), *pUBQ10::YFP-VAMP711 amiR vps3* (Takemoto et al., 2018). *vti11 pUBQ10::YFP-VAMP711* was obtained by crossing. Seeds were stratified at 4°C for 2 days in the dark and were grown on vertically orientated ½ strength Murashige and Skoog (MS) medium plates containing 1% sucrose under a long-day regime (16 h light/8 h dark) at 20–22°C.

### Chemicals

All chemicals were dissolved in dimethyl sulfoxide (DMSO) and applied in solid or liquid ½ MS medium. FM4-64 was obtained from Invitrogen/Thermo Fisher Scientific (MA, USA), propidium iodide (PI) and dexamethasone (DEX) from Sigma (MO, USA), 2′,7′-Bis(2-carboxyethyl)-5(6)-carboxyfluorescein acetoxymethyl ester (BCECF-AM) from Biotium (CA, USA), Wortmannin (WM) from MedChemExpress (NJ, USA) and Brefeldin A (BFA) from Alfa Aesar (MA, USA). VAC1 (N-[(2,4-Dibromo-5-hydroxyphenyl)methylideneamino]-2-phenylacetamide, ID 5326213), VAC1A1 (2-phenyl-N’-(2,4,6-tribromo-3-hydroxybenzylidene)acetohydrazide, ID 5326215), VAC1A2 (N’-(2-bromo-5-hydroxybenzylidene)-2-phenylacetohydrazide, ID 5326212), VAC1A3 (N’-(2,4-dibromo-5-hydroxybenzylidene)-2-(1-naphthyl)acetohydrazide, ID 5575792), VAC1A4 (N’-(2,4-dibromo-5-hydroxybenzylidene)-2-hydroxy-2-phenylacetohydrazide, ID 5568198) and VAC1A5 (N’-(3,5-dibromo-4-hydroxybenzylidene)-2-phenylacetohydrazide, ID 5326129) were obtained from ChemBridge/Hit2Lead (CA, USA) (https://www.hit2lead.com/search.asp?db=SC).

### VAC1

VAC1 used in this study was either obtained from ChemBridge/Hit2Lead (CA, USA) (https://www.hit2lead.com/search.asp?db=SC, ID 5326213) or synthesized in-house as described in VAC1 synthesis (see below). We performed LC-MS analysis to determine the similarity of synthesized and commercially available VAC1. HPLC chromatograms and MS spectra of the synthesized VAC1 were identical to the ones of the reference VAC1. The purity of the synthesized VAC1 (99.2 %) was higher than that of the purchased VAC1 (98.2 %). The IUPAC (International Union of Pure and Applied Chemistry) nomenclature was adapted to the VAC1 molecule (N-[(2,4-Dibromo-5-hydroxyphenyl)methylideneamino]-2-phenylacetamide) throughout the manuscript, however an alternative name for VAC1 (N’-(2,4-dibromo-5-hydroxybenzylidene)-2-phenylacetohydrazide) is used by the manufacturer Chembridge/Hit2Lead.

### VAC1 synthesis

**Scheme 1.**
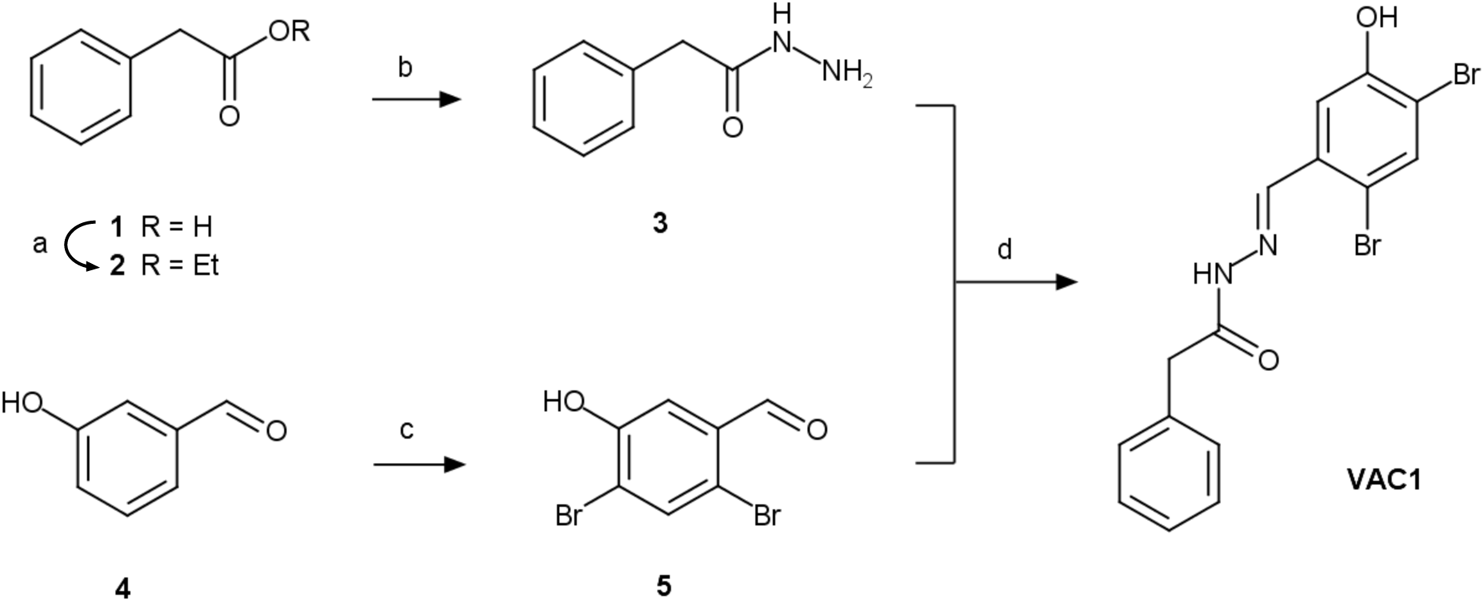
Preparative route to VAC1. Reaction conditions: a) EtOH, cat. Fe_2_(SO_4_)_3_/H_2_SO_4_, reflux, 3.5 h, 87 %; b) N_2_H_4_·nH_2_O (50-60% N_2_H_4_), MeOH, rt, 24-72 h, 68%; c) Br_2_ (2.12 eq.), DCM, rt, 22 h, 64 %; d) EtOH, reflux, 3 h, 76 %.

#### Experimental

Starting reagents 3-hydroxybenzaldehyde **4** and hydrazine hydrate were purchased from Sigma-Aldrich and phenylacetic acid **1** was purchased from Merck. These reagents were used as received without further purification.

Synthesis of **5** was accomplished by following the respective procedure in Kallmann et al., 2014 and **2** was prepared adapting Liang et al., 2004.

TLC was run on Silica 60 glass plates (Merck). Low-resolution mass-spectra were obtained on an Agilent 6340 Ion Trap instrument. For *m/z* values, the most abundant isotope of the isotope distribution is reported.

#### Ethyl phenylacetate (2)

A solution of **1** (4.08 g, 30 mmol) in abs. ethanol (150 ml) containing 50 mg of equimolar mixture of iron (III) sulfate and concentrated sulfuric acid was held at reflux for 3.5 h until no starting material was visible on TLC. Ethanol was distilled off and the residue was dissolved in DCM, washed with saturated aqueous solution of sodium bicarbonate, dried with anh. magnesium sulfate, stripped of solvent and purified by distillation in vacuo (bp 105-108°C at 13 Torr) to afford **2** as colorless liquid. Yield: 4.3 g (87%).

#### Phenylacetic acid hydrazide (3)

Hydrazine hydrate (2.81 ml, 45-54 mmol) was added dropwise to a solution of **2** (4.0 g, 24 mmol) in methanol (28 ml) and the mixture was agitated at rt for 24-72 h until complete consumption of the ester (TLC). Excess solvent was removed by distillation and, upon cooling, the residue crystallized as long thin needles. The mass was crushed, washed on a glass filter with cold water and recrystallized from 90% aq. ethanol. Yield: 3.0 g (82%) before and 2.5 g (68%) after recrystallization.

#### 2,4-Dibromo-5-hydroxybenzaldehyde (5)

To a solution of **4** (1.0 g, 8.2 mmol) in DCM (10 ml) was slowly added elemental bromine (2.78 g 17.4 mmol) and the reaction mixture was stirred at rt for 22 h. The reaction progress was monitored by HPLC. Upon completion, the reaction was quenched by dropwise addition of 15% aq. sodium thiosulfate (4.8 ml), stirred at rt for 1 h and filtered. The filter cake was washed with water (2×5 ml) and dissolved in acetic acid (6.4 ml) at 95°C. After cooling to 50°C, water (3.4 ml) was added dropwise to a stirred solution, resulting in crystallization of the product. The crystals were stirred for 4 h at 15°C, filtered, washed with water (2×5 ml) and dried. Yield: 1.68 g (73%). The product (pink crystals) was additionally purified by vacuum sublimation, affording **5** as white crystalline powder (1.48 g, 64%). *m/z* (ESI^−^): 278.8 [M-H]^−^.

#### N-[(2,4-Dibromo-5-hydroxyphenyl)methylideneamino]-2-phenylacetamide (VAC1)

A mixture of **3** (0.53 g, 3.53 mmol) and 1.02 eq of **5** in 9.6 ml of ethanol was held at reflux for 3 h until complete consumption of the starting material (TLC). Upon cooling, a copious white precipitate of the title compound was formed. It was filtered, washed with petroleum ether and dried. The final product was additionally purified by recrystallization from ethanol. Yield 1.1 g (76%). *m/z* (ESI^−^): 410.9 [M-H]^−^.

### UHPLC-UV-(SIR)MS

To determine the conversion of VAC1 into free PAA, 6-day-old seedlings were treated with DMSO or 10 µM VAC1 for 2.5 h in liquid medium and then flash-frozen in liquid nitrogen. The extraction and quantification methods for both analytes were optimized based on the previously described method (Pařízková et al., 2021). Briefly, the plant samples (∼30 mg FW) were extracted into 100% methanol and purified using liquid-liquid extraction method into 900 µL of extraction solution (methanol:H2O: hexane – 1:1:1), evaporated and dissolved in 50 µL of 100% methanol. 2 µL of each sample were injected onto the reversed-phase column (Kinetex C18 100A, length 50 mm, diameter 2.1 mm, particle size 1.7 μm; Phenomenex, CA, USA) and analysed by an Acquity UPLC H-Class System (Waters, Milford, MA, USA) coupled with an Acquity PDA detector (scanning range 190–400 nm with 1.2 nm resolution) and a single quadrupole mass spectrometer QDa MS (Waters MS Technologies, Manchester, UK) equipped with an electrospray interface (ESI). The analytes were eluted from the column within 9-min-linear gradient of 10:90 to 95:5 A:B using 0.1% acetic acid in methanol (A) and 0.1% acetic acid in water (B) as mobile phases at a flow rate of 0.5 ml.min-1 and column temperature of 40 °C. After every analysis, the column was washed with 95% methanol and then equilibrated to initial conditions (1.0 min). Both analytes were detected by Selected Ion Recording (SIR) using the positive and negative electrospray modes (ESI+ and ESI–) as follows: VAC1 detected as [M+H]+, m/z 411.0 and PAA as [M-H]-, m/z 135.0. The MS settings were optimized as follows: Source Temperature, 120 °C; Desolvation Temperature, 600 °C; Capillary Voltage, 0.8 kV. Chromatograms were processed by MassLynx V4.2 software (Waters) and quantification was performed from external calibration using a recovery factor. Additionally, M1-M3 metabolites of VAC1 were detected using a full-scan mode (m/z 50-1000) operated in ESI– with post-data-aquistion extraction of ion chromatograms for m/z 425.0, 571.0 and 613.0 for M1, M2 and M3, respectively. To determine the distribution of VAC1 metabolites, the UV chromatograms were extracted for λmax (291 nm) and the percentage representation of metabolites in plant extracts was determined by the integration of peak areas in respective retention times (M1, 4.37 min; M2, 5.26 min; M3, 5.39 min; VAC1, 6.65 min).

### UHPLC-HRMS

Samples of DMSO- and VAC1-treated plants were prepared as described above (UHPLC-SIR-MS). Plant extracts (2 µL) were injected onto a Kinetex C18 column and separated using chromatographic conditions as described above. The high-resoluion mass spectrometry (HRMS) analysis was achieved by a hybrid Q-TOF tandem mass spectrometer Synapt G2-Si (Waters MS Technologies) as described previously (Buček et al., 2018). Briefly, the effluents were introduced into the HRMS instrument (ESI–; Capillary Voltage, 0.75 kV; Source Offset, 30 V; Desolvation/Source Temperature, 550/120°C; Desolvation/Cone Gas Flow, 1000/50 l hr−1; LM/HM Resolution, 2.8/14.75; Ion Energy 1/2, 0.5/1.0 V; Entrance/Exit Voltages, 0.5 V; Collision energy, 6 eV). The determination of exact mass was performed by the external calibration using lock spray technology and a mixture of leucine/encephalin (1 ng.μl-1) in an acetonitrile and water (1 : 1) solution with 0.1% formic acid as a reference. Data acquisition was performed in full-scan mode (50–1000 Da) with a scan time of 0.5 s and all analytes were detected as [M-H]– in the MS spectrum. All data were processed using MassLynx 4.1 software (Waters). The accurate masses of VAC1 metabolites were calculated and then used to determine the elemental composition of the analytes with a fidelity ranging from 1.6 to 2.6 ppm. The VAC1 metabolites were identified based on correlation of the theoretical monoisotopic weights of deprotonated forms [M-H]- and detected accurate masses of each precursor of M1-M3 as well as based on the presence of two bromine stable isotopes (79Br and 81Br) in HRMS spectrum resulting the characteristic isotope pattern of dibromo derivatives.

### Phenotype analysis

Vacuolar occupancy and vacuole surface area as well as cell surface area and BFA body volume was quantified in 6 days old seedlings. For 3D reconstructions of cells, confocal images were processed using Imaris (vacuolar occupancy of cells, BFA bodies, vacuole and cell surface) as described previously (Dünser et al., 2019). BCECF staining was performed as described previously (Scheuring et al, 2015). For the analysis of main root growth, 6-day-old seedlings were used, unless indicated otherwise. For the analysis of main root growth of amiR *vps3* and amiR *vps39* seedlings were grown for 4 days before transfer to DMSO, DEX, VAC1 or DEX + VAC1 and then grown for additional 3 days. Plates were scanned and root length assessed using ImageJ. Selection of pUBQ10::YFP-VAMP711 amiR *vps3* seedlings for confocal microscopy analysis was done as described previously (Takemoto et al., 2018).

### Confocal Microscopy

For image acquisition a Leica TCS SP5 (DM6000 CS) or a Leica SP8 (DMi8) confocal laser-scanning microscope, equipped with a Leica HCX PL APO CS 63 × 1.20 water-immersion objective, was used. GFP and BCECF were excited at 488 nm (fluorescence emission: 500–550 nm), YFP and FM4-64 at 514 nm (fluorescence emission YFP: 525–578 nm; fluorescence emission FM4-64: 670-790 nm), and PI at 561 nm (fluorescence emission: 644–753 nm). Roots were mounted in PI solution (0.02 mg/mL) for the counterstaining of cell walls. Z-stacks were recorded with a step size of 420 nm (for 3D reconstructions) or 1.5 µm (for maximum projections of VAC1 bodies).

### 3-D reconstruction of cells

Imaris 8.4.0 and 9.0 were used for the reconstruction of cell, vacuole and BFA body volumes. Based on the PI channel, every 3rd slice of the z-stack was utilised to define the cell borders using the isoline, magic wand or manual (distance) drawing functions in the manual surface creation tool. After creating the surface corresponding to the entire cell, a masked channel (based on BCECF or FM4-64) was generated by setting the voxels outside the surface to 0. Subsequently, a second surface (based on the masked BCECF or FM4-64channel) was generated automatically with the smooth option checked. Surface detail was set to identical values within each region (early and late meristem and early and late elongation zone) to ensure comparability of obtained surface area values. The obtained surface was visually compared to the underlying BCECF or FM4-64 channel and, if necessary, the surface was fitted to the underlying signal by adjusting the absolute intensity threshold slider. Finally, volumes and surface areas of both surfaces were extracted from the statistics window.

### Small molecule screen

Roughly 20 pUBQ10::YFP-VAMP711 seeds were germinated in 12 well plates with each well containing 1.5 ml liquid ½ MS+ supplemented with solvent control (DMSO) or 50 µM of small molecules. 4-day-old seedlings were screened for enhanced fluorescence signal using a fluorescence binocular. 12 hit compounds were further examined using confocal microscopy (see Suppl. Fig. 1 A, B). The small molecules used for the primary screen were identified as inhibitors of tobacco pollen germination (Drakakaki et al., 2011).

### Quantification of VAC1- and BFA bodies

Maximum z-projections of 3 – 8 slices (step size 1.5 µm) and a ROI of 5 µm x 5 µm were used for quantification of the FM4-64 fluorescence signal. 4 – 10 roots were quantified with 3 bodies per root.

### Colocalization analysis

Quantification of colocalization was done using ImageJ. 2 – 3 atrichoblast cells in the specified regions were used per root. The area was marked and cropped (Image > Crop), subsequently the surrounding of the cell was cleared (Edit > Clear outside) and the channels were split (Image > Color > Split channels). Colocalization (Analyze > Colocalization > Colocalization threshold) was analyzed and the values for Rtotal as well as the scatter plots were extracted.

### Time course experiments

Seedlings were mounted on agar blocks containing solvent control DMSO or VAC1. The agar blocks were placed in chamber slides and subsequently installed on an inverted confocal microscope (Leica SP8). Z-stacks (3 – 5 slices, step size: 1.5 µm) were acquired every 10 minutes and maximum z-projections used for quantification of root tip displacement and cell length.

### Cell length analysis

6-day-old seedlings were used for cell length measurements of differentiated atrichoblasts (Fig. 1 C, D). Fully differentiated cells were identified as described previously (Löfke et al., 2015).

## Acknowledgements

We are grateful to N. Geldner, S. Robert, K. Schumacher and T. Ueda for sharing published material. The library of 360 bioactive small molecules was kindly provided by N. Raikhel. We thank the BOKU-VIBT Imaging Center and the ZBSA-Life Imaging Center for access.

This work was supported by the Ministry of Education, Youth and Sports of the Czech Republic (European Regional Development Fund-Project “Plants as a tool for sustainable global development” No. CZ.02.1.01/0.0/0.0/16_019/0000827 to O. N.), The Austrian Academy of Sciences (ÖAW) (DOC fellowship to K.D.), Austrian science fund (FWF; P 33044 to J.K.-V.), German Science fund (DFG; 470007283 and CIBSS – EXC-2189 to J.K.-V.), and the European Research Council (ERC; 639478-AuxinER to J.K.-V.).

## Author contributions

KD, CL and JK-V designed research. KD, MS, CL and NX conducted experiments. BP and ON performed UHPLC–SIR–MS and UHPLC-HRMS. SM synthesized VAC1. All authors analyzed data. KD and JK-V wrote the manuscript and all co-authors commented on the manuscript.

## Conflict of interest

The authors declare that they have no conflict of interest.

## Supplemental Figure legends

**Supplemental Figure 1.**
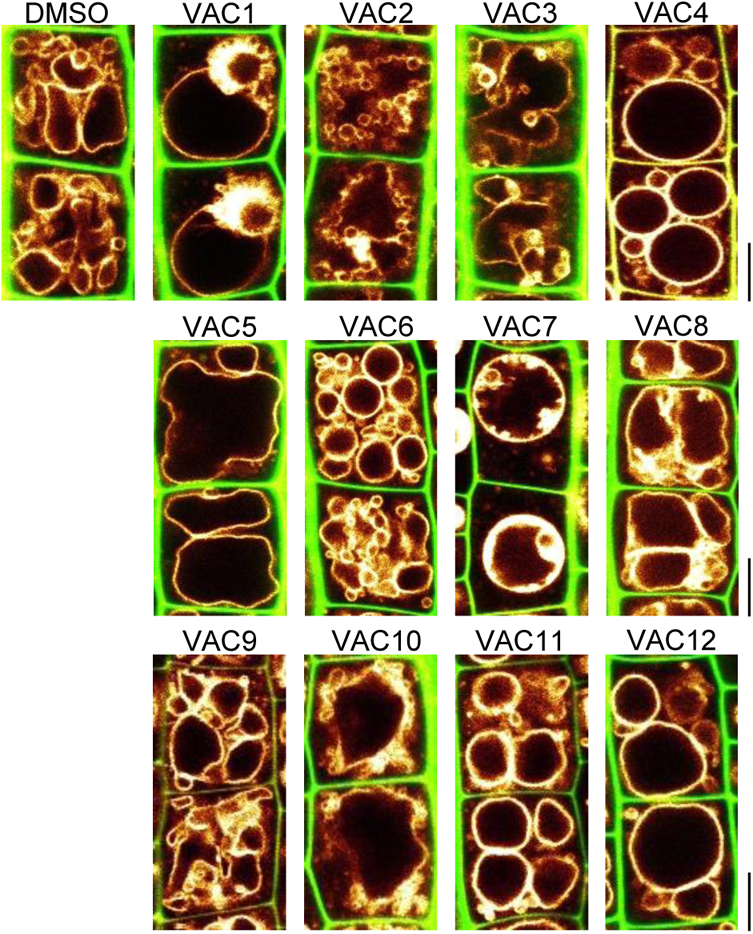
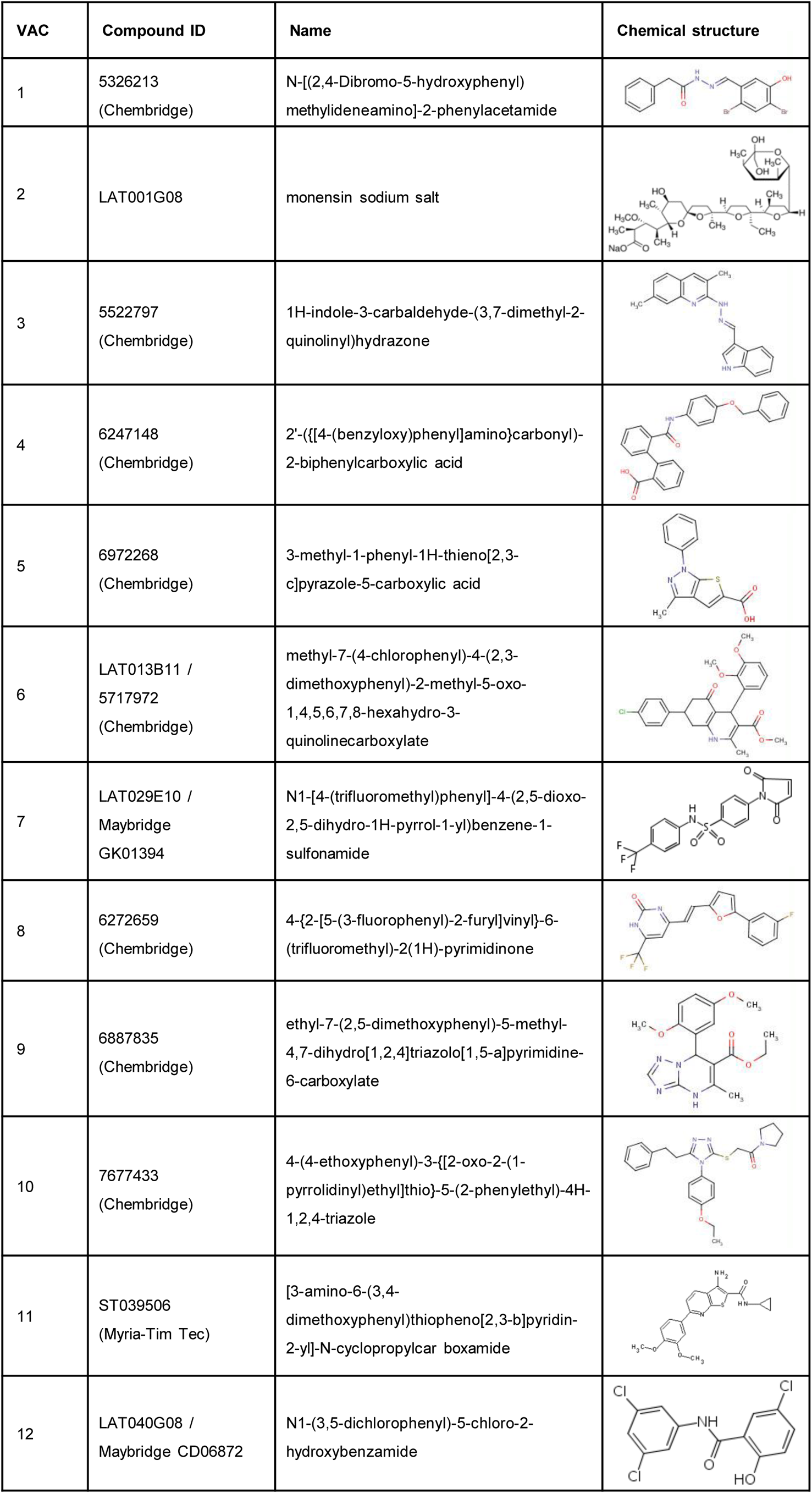
Bioactive small molecules impact vacuolar morphology. (A) Cell wall and vacuolar membrane in late meristematic atrichoblast cells were visualized with cell wall stain propidium iodide (PI) (green) and pUBQ10::YFP-VAMP711 (yellow). 6-day-old seedlings were treated with solvent control (DMSO) or VAC1 (7.5 µM, 1 h), VAC2 (7.5 µM, 5 h), VAC3 (5 µM, 5 h), VAC4 (23.3 µM, 5 h), VAC5 (10 µM, 5 h), VAC6 (50 µM, 5 h), VAC7 (7.5 µM, 1 h), VAC8 (7.5 µM, 2.5 h), VAC9 (100 µM, 5 h), VAC10 (100 µM, 2.5 h), VAC11 (25 µM, 5 h), VAC12 (100 µM, 6 h) in liquid medium. Scale bars: 5 µm. (B) Table shows compound IDs, full name and chemical structure.

**Supplemental Figure 2.**
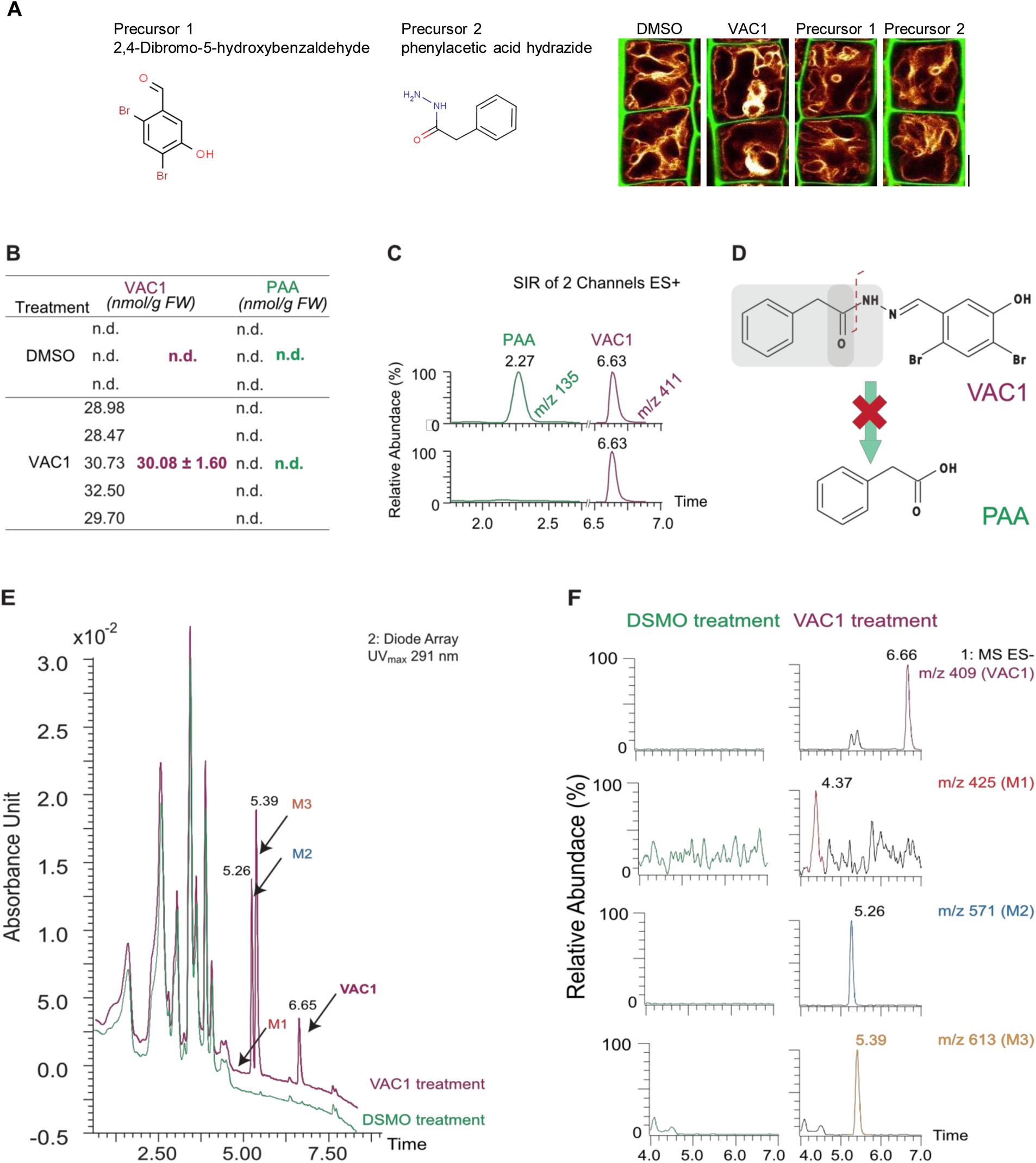
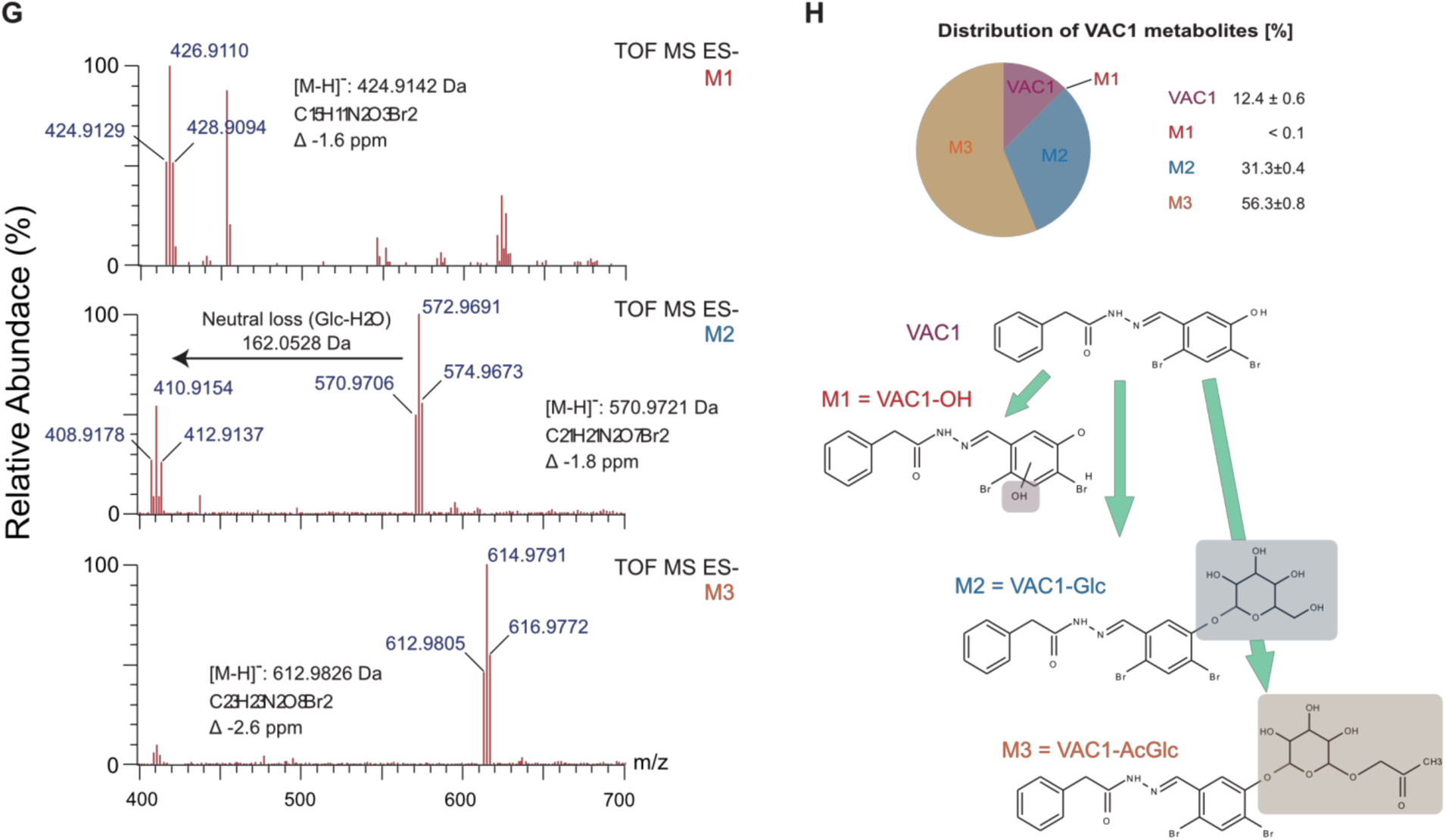
Characterization of VAC1 stability *in planta*. (A) Cell wall and vacuolar membrane in late meristematic atrichoblast cells were visualized with PI (green) and pUBQ10::YFP-VAMP711 (yellow). 6-day-old seedlings were treated with solvent control (DMSO) or 10 µM VAC1, 10 µM Precursor 1 or 10 µM Precursor 2, respectively, in liquid medium. Scale bar: 5 µm. (B) 6-day-old seedlings were treated with DMSO or 10 µM VAC1 for 2.5 h in liquid medium, flash-frozen in liquid nitrogen and subsequently extracted by liquid-liquid extraction (Pařízková et al., 2021). Quantification of VAC1 and PAA levels was performed by UHPLC-UV-MS system operating in selected ion recording (SIR) mode and calculated from an external calibration using a recovery factor. No PAA was detected in DMSO- and VAC1-treated seedlings. Values are means ± SD, n = 3 for DMSO and n = 5 for VAC1 treatment. (C) Representative SIR chromatograms of PAA and VAC1 comparing retention times obtained by measuring a standard mixture (top ion chromatogram) with analysis of plant extract (bottom ion chromatogram). (D) The potential of VAC1 to be metabolized into free auxin PAA (green rectangle) by cleavage of the peptide bond (red rectangle) was not confirmed. (E) UV chromatograms (λ_max_ = 291 nm) of DMSO- and VAC1-treated plant extracts obtained in (B) revealed peaks, additional to VAC1 (retention time: 6.65 min), corresponding to three unidentified metabolites M1, M2 and M3 eluting from the LC column at 4.37 min, 5.26 min and 5.39 min, respectively. (F) Representative extracted-ion chromatograms of VAC1 and M1-M3 metabolites with their respective retention times and precursor masses were obtained by UHPLC-UV-MS analysis in negative full scan mode from DMSO- and VAC1-treated plant extracts. (G) HRMS spectra of three VAC1 metabolites M1-M3. All metabolites were identified using an UHPLC-HRMS method based on correlation of the theoretical monoisotopic weights of deprotonated forms [M-H]^−^ and detected exact masses of each precursor of M1-M3 as well as based on the presence of two bromine stable isotopes (^79^Br and ^81^Br) in VAC1 structure showing the characteristic isotope pattern of dibromo derivatives. (H) Predicted chemical structures of VAC1 metabolites and their distribution (%) in extracts of 6-day-old seedlings treated with 10 µM VAC1 for 2.5 h in liquid medium. The percentage representation of metabolites was determined by the integration of respective peaks in the UHPLC-UV chromatograms (λ_max_ = 291 nm). Values are means ± SD (n = 5). The VAC1 metabolites were identified as VAC1-OH (M1), VAC1-Glc (M2) and VAC1-AcGlc (M3). The color rectangles display the metabolite changes in VAC1 structure (hydroxylation/glycosylation) in the presumable positions.

**Supplemental Figure 3.**
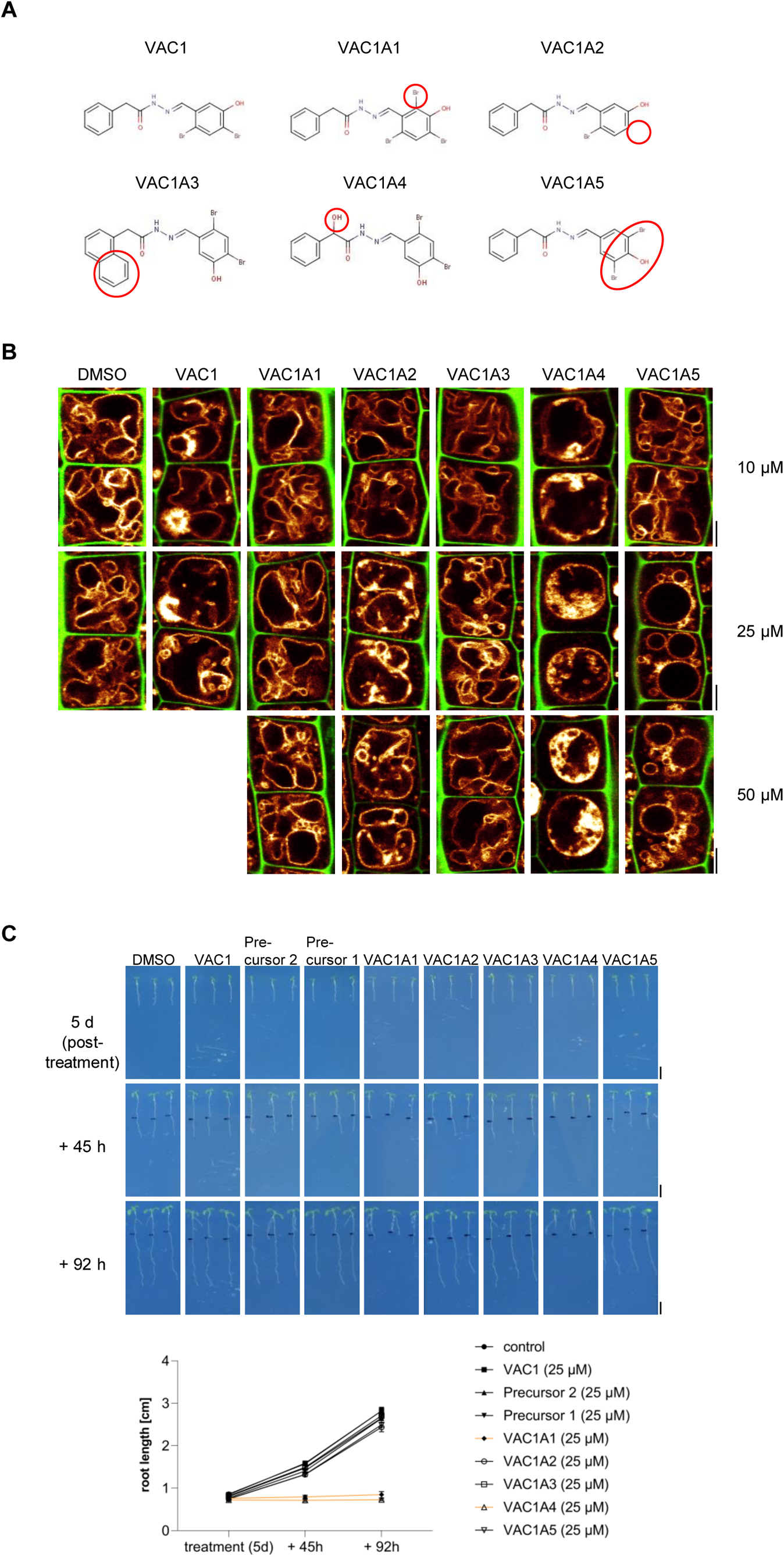
Characterization of VAC1-derivatives. (A) Structural formulas of the examined VAC1-derivatives (VAC1As). (B) Cell wall and vacuolar membrane in late meristematic atrichoblast cells were visualized with PI (green) and pUBQ10::YFP-VAMP711 (yellow). 6-day-old seedlings were treated with solvent control (DMSO) or indicated concentrations of VAC1 and VAC1As, respectively, for 2.5 h in liquid medium. Scale bar: 5 µm. (C) Representative images and quantification of main root length after compound treatment. 5-day-old seedlings were treated with 25 µM DMSO, VAC1, Precursors and VAC1As, respectively, for 2.5 h in liquid medium. The seedlings were then transferred to agar plates and scanned at the timepoints indicated. Scale bars: 0.5 cm. Graph shows average root length values with s.e.m. at indicated timepoints.

**Supplemental Figure 4.**
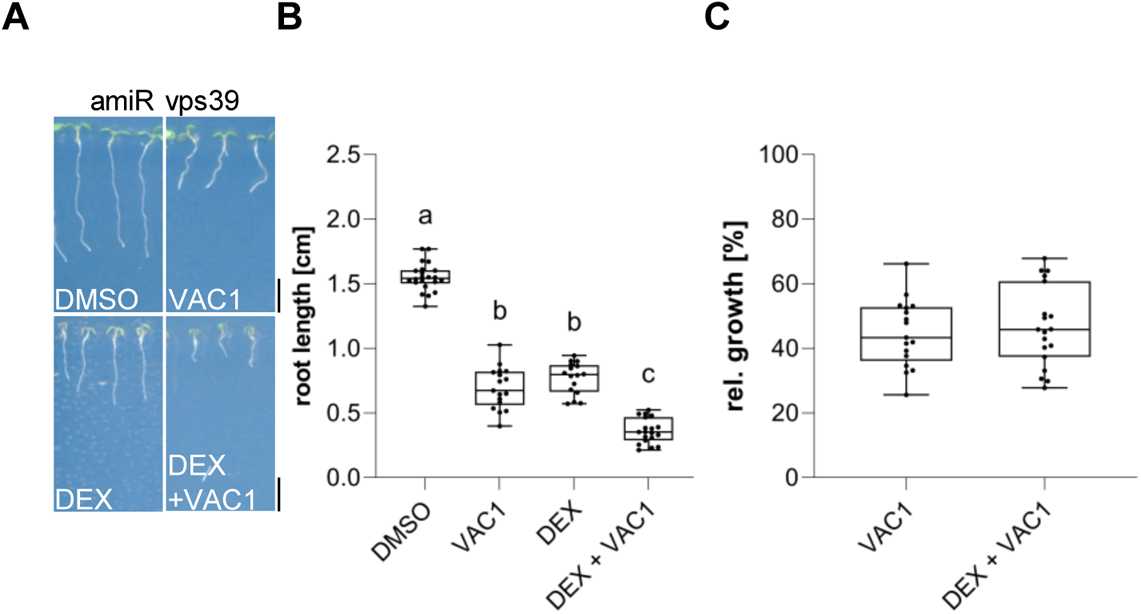
VPS39-deprived roots are sensitive to VAC1. (A) Representative images (scale bars: 0.5 cm) and (B) quantification of main root length of 7-day-old amiR *vps3* seedlings germinated on solvent control medium (DMSO, n = 22), 10 µM VAC1 (n = 17), 30 µM DEX (n = 16) or 10 µM VAC1 and 30 µM DEX (n = 19), respectively. One-way ANOVA with Tukeýs multiple comparisons test (b: P < 0.0001, c: P < 0.0001). (C) Boxplots show relative growth of VAC1 samples (compared to DMSO) and DEX + VAC1 samples (compared to DEX). Student’s t-test (ns). Boxplots: Box limits represent 25^th^ percentile and 75^th^ percentile; horizontal line represents median. Whiskers display min. to max. values. Data points are individual measured values. Representative experiments are shown.

**Supplemental Figure 5.**
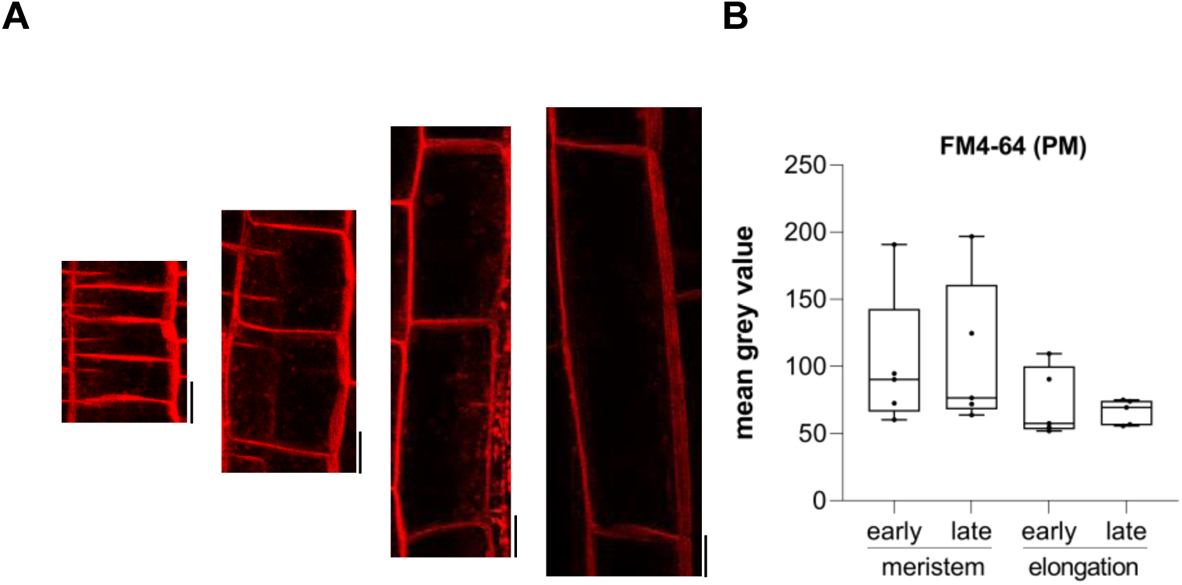
FM4-64 evenly stains transversal plasma membranes in the meristem and elongation zone. (A) Representative images (scale bars: 10 µm) of 6-day-old Col-0 seedlings that were pulse stained with 4 µM FM4-54 (red) for 5 min in liquid medium before image acquisition. (B) quantification of FM4-64 fluorescence signal of transversal plasma membranes in the early (n = 5) and late (n = 5) meristem and in the early (n = 5) and late (n = 5) elongation zone. One-way ANOVA with Tukeýs multiple comparisons test (ns). Boxplots: Box limits represent 25^th^ percentile and 75^th^ percentile; horizontal line represents median. Whiskers display min. to max. values. Data points are individual measured values. Representative experiments are shown.

## Notes

### Competing Interest Statement

The authors have declared no competing interest.

